# Beyond the metropolis: Street tree densities and resident perceptions on ecosystem services in small urban centers in India

**DOI:** 10.1101/2022.09.28.509699

**Authors:** Krishna Anujan, Nandini Velho, Giby Kuriakose, Ebin P J, Vivek Pandi, Harini Nagendra

**Affiliations:** Columbia University in the City of New York; Srishti Manipal School of Art, Design and Technology, Bengaluru; Sacred Heart College, Kochi; Manipal Centre for Natural Sciences, Manipal Academy of Higher Education, Manipal; Azim Premji University, Bengaluru

**Keywords:** urban sustainability, small cities, ecosystem services, street trees

## Abstract

The role of urban street trees has been extensively studied in large metropolises, where they contribute significantly to faunal habitat, provide critical ecosystem services to residents and contribute to human well-being. On the other hand, rapidly urbanizing cities in India have been poorly studied, despite multiple types of irreplaceable losses related to tree cover. However, being early in their urbanization history, these centers also represent opportunity for urban sustainability with potentially high remnant vegetation and human-nature connections. While megacities in developed countries move towards biophilic urban planning and participatory decision making, basic information on tree communities and their perceived services is a bottleneck in achieving these goals in such small urban centers. We assessed the street tree community and resident perceptions of ecosystem service values in Kochi and Panjim, two coastal cities in India under rapid development, through a combination of field measurements (258 transects, 931 trees) and semi-structured interviews (497 individuals). We found that mean street tree density is low in both cities, especially so in Kochi, and corresponds to perceptions of recent change in tree cover (−28% in Kochi, −11% in Panjim). The street tree community in both cities were dominated by ornamental avenue trees such as *Albizia saman* and *Peltophorum pterocarpum*, but native coastal species like *Cocos nucifera*, *Terminalia catappa* and *Thespesia populnea* were also common. Despite recent urban growth, residents in both cities reported low value of trees for food, fodder and medicine, but high value for regulating services like shade and water. Moreover, we found strong evidence for aesthetic and cultural values of trees in both cities, including through qualitative interviews. Our study establishes critical baselines for biophilic planning in these small urban centers towards urban sustainability.

## Introduction

Urban street trees and greenery are important and effective solutions to improve sustainability and resilience of cities as well as the well-being of residents (Beatley & Newman, 2013; Blaustein, 2013; T. A. Endreny, 2018). Trees contribute directly to biodiversity conservation and provide critical patch connectivity for plants and animals (Henry et al., 2017; Jaganmohan et al., 2018; Somme et al., 2016; Wood & Esaian, 2020). Further, urban trees reduce pollution, contribute to climate resilience by mitigating stormwater and reducing urban heat island effects as well as improve physical and mental health of residents (Beyer et al., 2014; Bratman et al., 2019; Parker & Simpson, 2020; Snep et al., 2020; Triguero-Mas et al., 2015; Willis & Petrokofsky, 2017). Urban trees could save megacities nearly $500 million annually in public health, stormwater management and climate mitigation (T. Endreny et al., 2017). Recognising the value of biophilic cities, urban planning in major cities is shifting towards regreening and rewilding (Beatley & Newman, 2013; Blaustein, 2013; Lehmann, 2021). Street trees could be a low-cost tool in creating sustainable cities in developing countries, although other solutions (eg: green roofs) are gaining popularity in the developed world (Snep et al., 2020). Efforts at protecting street trees and urban habitats make significantly high contributions to five UN Sustainable Development Goals, namely zero hunger, good health and well-being, clean water and sanitation, sustainable cities and communities and climate action (T. A. Endreny, 2018; Food and Agriculture Organization of the United Nations, 2016). To accurately assess the sustainability of cities, integrative approaches need to characterize nature in the city as well as human-nature connections.

Urban ecology and sustainability is increasingly considered in planning, but research is biased towards larger metropolises in the global north. Although 41% of the global urban population lives in smaller urban centers, most studies on urban nature are focussed on large, developed, metropolises and fail to represent this population (Kendal et al., 2020). Moreover, while urbanization is presently correlated with countries in the global south (Jain & Korzhenevych, 2020), research and policy on urbanization is influenced by the north (Nagendra et al 2018). India, one of the fastest growing economies in the world, has been relatively slow to wake up to the opportunities and challenges of urbanization occurring at multiple scales (Sankhe et al., 2011). Growth trajectories of cities in the global south are different from global north realities. In India, there has been a rise of smaller “census” towns, making a complex rural-urban matrix. However, as per the last population census, 30% of the country’s urban growth has taken place in these census towns - smaller urban centers that are governed by village bodies (Tripathi & Mitra, 2022).This ‘shock urbanization’ in India is leading to disentanglement from infrastructure services and physical and social short-comings (Jain & Korzhenevych, 2020; Rode et al., 2008). Emerging studies from larger metropolises such as Bengaluru in India, which saw rapid infrastructure in the last decade with phasing out of large-sized trees in older green spaces (Gopal et al., 2015; Nagendra & Gopal, 2010). Being in the early stages of urbanization, smaller urban centers can potentially have higher green cover and more native species, characteristic of rural landscapes, presenting an opportunity to proactively plan for “biophilic” cities.

Fostering biophilic urbanism works when people know and care about trees and their value along with establishing ecological baselines (Beatley & Newman, 2013). Street trees provide a variety of ecosystem services and health benefits to residents such as food, fodder, firewood, regulating services related to temperature, water, air quality and cultural services that are aesthetic, spiritual and place-based (Salmond et al., 2016). Besides the positive effects of reducing atmospheric carbon dioxide, urban trees reduce particulate matter and urban heat island effects (Sinha et al., 2021; Vailshery et al., 2013; Willis & Petrokofsky, 2017). Tree lined avenues can support eco-friendly behaviors such as walking and cycling (Lusk et al., 2020). More broadly, urban neighborhoods with trees are often associated with better mental and physical health (Beyer et al., 2014; Taylor et al., 2015; Triguero-Mas et al., 2015). However, street trees are multifunctional and there are often trade-offs between their services and risks such as damage to property and infrastructure, pollen load and associated allergies, where positive perceptions of trees co-exist with “Not-In-My-Backyard” attitudes (C. O. Fernandes et al., 2019). Age, gender and education correlate with varying perceptions of cost-benefit trade-offs, assessments and/or indifference towards ecosystem services (C. O. Fernandes et al., 2019; Graça et al., 2018; Kloster et al., 2021). We know little about how residents in smaller cities especially in the global south experiencing urbanization perceive these diverse ecosystem services and their values. In these areas where trees were more abundant and accessible in the recent past, it is likely that provisioning services like food and medicine as well as aesthetic, religious and cultural values may be perceived to be important.To maximize the benefits of street trees to residents and to create biophilic cities, it is important to understand the changing ecological context of the cities along with their perceived values (Dobbs et al., 2018; Lehmann, 2021).

Panjim and Kochi are coastal cities in India that are growing in different ways with a past rich in natural and cultural heritage. Both located along the Arabian Sea coast, these cities have been multicultural trade centers for centuries, playing prominent roles in historical trade with Europe, the Middle-East and many African countries. As of the 2011 census, Kochi is a Tier-II city, with a population of over 6 lakh while Panjim is a Tier-III city with population less than 1 lakh. They have a growing trend in infrastructure, as part of the National Smart Cities Mission, an urban renewal and retrofitting programme, and in population, and are likely to be reclassified into higher tiers in the 2021 census. These cities are also frequented by tourists - from riverine backwaters to built heritage, outlining a strong need for prioritizing biodiversity in urban planning. Thus there is a timely opportunity to balance urban planning with preserving green spaces, collaboratively through multiple stakeholder engagement, setting an example for other growing cities in India. A first step towards these is assessing the status of green spaces and residents’ perspectives about them.

We assessed the abundance, distribution and diversity of street trees in Kochi and Panjim and interviewed residents to understand both status and perceptions of green spaces in these cities. Specifically, we aimed to (i) document present-day street tree abundance (ii) understand the distribution of trees in terms of species and size for setting ecological baselines (iii) and document perceptions, personal and cultural associations and changes in the city’s greenery through residents’ lived experience. We intend for this study to establish critical baselines for continuing work on documenting urban green spaces, their use and access in small, growing urban centers in the region in the future.

## Methods

Kochi (9.97°N 76.28°E) and Panjim (15°29′56′′N 73°49′40′′E) are coastal cities at sea level with a tropical monsoon climate and close to the Western Ghats biodiversity hotspot. An inventory of biodiversity by the State Biodiversity Board of Goa in Panaji in 2014 documented over 200 faunal species (Goa State Biodiversity Board & Corporation of the City of Panaji, 2014). Although the City Biodiversity Index prepared by the Kochi Municipal Corporation (Kochi Municipal Corporation, 2020) only documented 50 faunal species, a more intensive localized faunal survey, Joseliph & Davis, 2014, documented 193 species - 64 invertebrates and 129 vertebrates, including aquatic species. The urban area in Kochi has increased about five-fold in the last three decades and the land use pattern has changed dramatically (Abhimanyu et al., 2020). Panjim’s city’s population and growth trend over the past decades is unclear due to city ward maps not being publicly available and the change in the administrative area, but there has been an urban population clustering to Panjim’s municipal area and to adjacent village areas (Capacity Building for Urban Development Project, 2015). We conducted field studies in Kochi and Panjim on street trees and residents’ perceptions of green spaces in their cities from August 2019 to February 2020. We collected field data with the research assistance of students from local universities and colleges who were trained in data collection protocols across sites.

### Sampling design

We conducted our study within the administrative boundaries of the Kochi Municipal Corporation in Kochi and within the Assembly Constituency of Panaji as well as the adjacent constituency of Taleigao due to the smaller size of Panaji and the relative similarity between these areas. We used boundary files of Kochi and Panjim downloaded from public sources (Panjim and Taleigao derived using map from https://www.diva-gis.org/gdata and Kochi derived using map from https://projects.datameet.org/indian_village_boundaries/). Using QGIS, we overlaid regular 0.5 km x 0.5 km grids on both these cities. Our grid size was the same for both cities, but given the larger extent of Kochi we sub-sampled grids. For both cities in each grid we generated 3 random locations as transect start points of which we picked two on field. We sampled 258 transects; 106 in Panaji (52 in Panaji and 54 in Taleigao) and 152 in Kochi. Some areas in Kochi were excluded from the sample that belonged to the Indian National Army and were not accessible. For our analyses, we consider Panaji and Taleigao as “Panjim” together because of overlapping confidence intervals of tree densities and interviewee profiles (except education). We recorded and collated transect measurements, tree measurements and resident interviews using Epicollect5 (Aanensen et al., 2014). All statistical analyses were then performed using R version 4.1.3 and RStudio (R Core Team, 2022). Data configuration was done using *plyr* and *reshape2* (Wickham, 2020a, 2020b). For all variables measured in the study, data reported was bootstrapped using packages *boot* and *confintr;* all bootstrapping was performed 9999 times using the ci_mean function and the mean estimate and 95% confidence interval derived(Canty & Ripley, 2021; Mayer, 2022). Figures were made using the packages *ggplot2, ggcorplot*, *gridExtra, sp, rgdal, rgeos* and *prettymapr* (Auguie, 2017; Bivand et al., 2022; Bivand & Rundel, 2021; Dunnington, 2017; Pebesma & Bivand, 2021; Wickham, 2016).

### Characterizing the tree community

From each selected transect start point, we walked transects of 100 m length along the road in a direction where road width stayed constant and there was no overlap with adjacent transects, to characterize the tree community. In situations where road width was similar in both directions and there was no overlap, the transect direction was chosen randomly. At the start and end of every transect we measured road-width using a Tacklife HD60 laser distance meter. The average road width sampled was 7.00 m (95% CI = 6.40, 7.65) for Panjim and 5.63 m (95% CI = 5.11, 6.21) for Kochi. We measured and identified all trees and palms >10cm girth at breast height (gbh) on either side of the road that were in public property. For highways, we considered trees that were on either side of the road as well as on the median.

For each tree encountered during the transect surveys, we recorded species, gbh, height and environment around the tree base. We identified the species of the tree to the lowest possible taxonomic level using field guides like Trees of Delhi (Kishen, 2005). In case of unknown or unclear IDs, we took photographs and/or samples of the species which were identified later. To understand the nature of the built environment around the tree, we recorded whether or not the tree base was surrounded by cement, concrete or any other building material. We measured the height of the tree using the laser distance meter. We measured three points on the tree - the distance to the highest point (h), the horizontal distance (d) to the trunk and the distance to the base (b) of the tree. We also recorded a visual estimate (v) to add or subtract where the highest point was not tracked by laser and the chin height (t) of the observer. We calculated the height of the tree using these values differently for trees leaning towards and away from the observer with separate basic geometric equations.

Leaning toward:

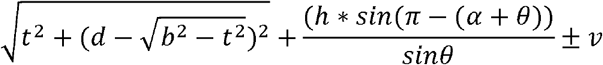

Where

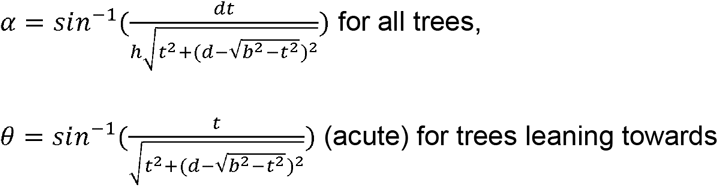

And

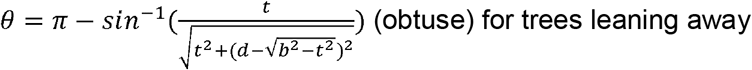

We measured the girth at breast height for every separate stem at 1.3 m from the ground using a standard tailor tape with an accuracy of 1 mm. We calculated the basal area for each tree by calculating the basal area of each stem as 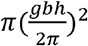 and summing across all stems. As there were some discrepancies with the gbh measurements and some species identification, we conducted a round of data validation in 2021 and excluded 10 transects (17 trees) from the basal area analysis.

Using the tree community measurements, we calculated Simpson’s diversity and Shannon diversity at the city level and at the transect level. For a community with *N* individuals across *i* species, each with abundance *n*_*i*_, we calculated relative abundance *p*_*i*_ as 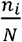. Simpson’s diversity was then calculated as Σ_*i*_ *p*_*i*_^2^ and Shannon diversity as Σ_*i*_ *p*_*i*_*log*_*e*_*p*_*i*_ We characterized the species composition in both cities by calculating, for each species, the relative abundance (number of individuals of species/total number of individuals), relative dominance (basal area of species/total basal area), relative frequency (proportion of transects in which the species occurred) and species importance (relative abundance + relative dominance + relative frequency).

### Documenting residents’ perceptions

At the start and end of each transect, we interviewed one respondent who was willing to speak given that there was a place (eg: house or shop) or a person who was willing to speak at the transect. We used a combination of semi-structured questionnaires and open-ended questions to understand human-tree relationships in Panjim and Kochi and specifically asked how citizens perceive their green spaces and biodiversity. We interviewed a total of 314 respondents in Kochi and 183 in Panjim. In Kochi, 152 identified as female, 162 as male while in Panjim, 76 identified as female, 107 as male. The mean age of respondents was 48.3 years in Kochi (95% confidence interval: 44.5, 53.4) and 43.9 years in Panjim (95% CI: 41.4, 46.4). Respondents had spent an average of 27.5 years (95% CI: 25.4, 29.5) living in Kochi and 27.8 years (95% CI: 24.9, 30.7) in Panjim. In Kochi, respondents were formally educated for an average of 11.6 years (95% CI: 11.3, 12.0) while in Panjim it was 10.5 years (95% CI: 9.8, 11.2). There was a significant difference in the number of years of formal education between respondents in Panjim (11.30; 95% CI: 10.50, 12.10) and Taleigao (9.24; 95% CI: 8.07, 10.40), but the interviewee profile was similar in the other two aspects.

Our interview questionnaire assessed the status, use and ecosystem services of urban trees. All interviews were conducted face-to-face and lasted approximately 45 minutes. The questionnaire was translated into Malayalam and Konkani for Kochi and Panjim respectively and conducted in these languages when necessary. The questionnaire was divided into sections that assessed (i) perceptions of change, and (ii) uses, beliefs, values and services of trees in their area. For all analyses, we considered responses where the respondent was neither “unsure” nor “uncomfortable” and excluded this data.

We asked questions about 14 different potential ecosystem services provided by trees and asked respondents to rank each specific use in their area. Depending on the service, respondents rank it either on a four-point scale of frequency of use - “frequently”, “occasionally”, “rarely” or “never” - or on popularity of use “everyone does”, “quite a few do”, “very few”, “not at all”. For analysis, both of these scales were interpreted as “very high value”, “some value”, “little value” and “no reported value”, allowing all services to be analyzed similarly. For each service, to understand the range of perceptions, we calculated probabilities of reporting “very high” value and “no reported value” in each city. To understand perceptions of multiple values and services provided by trees, we combined the reported ecosystem services into categories following the Millenium Ecosystem Assessment report (Alcamo et al., 2003). Specifically, we grouped them into (a) provisioning services: harvest for food, livestock fodder, medicine, timber or firewood, (b) regulating services: shade for buildings and roads, shade for farming, water regulation or (c) aesthetic, cultural and religious services: demarcating boundaries, beauty, connection with spirits, associated with religious buildings, associated with other cultural buildings. For each respondent, we counted the number of services in each category reported as occurring with “very high” value in their area. To understand whether the perceptions of these services, either individually or within their categories, were correlated among respondents, we explored correlations between the reporting of “very high” value.

We asked respondents when they had last been to a park or a green space and recorded a qualitative response. We analyzed this data by searching for specific terms in the responses to categorize frequency of visit - responses that mentioned “day”, “everyday” or “daily” were considered “within the week”, responses with the string “week” were considered as “within the month” and responses with “month” were considered “within the year”. All responses without any of these terms were considered as “longer than a year”. We summarized the frequency of these responses in both cities. Further, we asked respondents about tree cutting and planting activities in their area, specifically whether it occurred “frequently”, “occasionally”, “rarely” or “never”. Excluding “unsure” or “uncomfortable” responses, we calculated bootstrapped mean and confidence intervals in each city for probability of tree cutting and planting in each of the categories. We also identified perceived trends in tree abundance in the last decade by asking respondents to quantify change in tree abundance in their area as 25 to 50% decrease, 5-25% decrease, no change, 5-25% increase, 25-50% increase). We analyzed this information by assigning mid-value to each interval (25 to 50% decrease: −37.5, 5-25% decrease: −10 no change: 0, 5-25% increase: 10, 25-50% increase: 37.5) and assessing the mean perception of change. For this analysis, we only considered responses of interviewees who had lived in the city for more than 10 years.

## Results

### Tree community

Panjim ranked higher than Kochi in roadside tree abundance and diversity. At the transect level as well, Panjim had significantly higher values of tree density, species richness, Simpson and Shannon diversity (Table 1). Out of a total of 913 trees, the mean number of trees in a transect in Kochi was 1.54 (95% CI: 0.95, 2.26, n= 234) and in Panjim was 6.41 (95% CI: 5.26, 7.64, n = 679). The maximum number of trees in a transect in each city was comparable; 27 in Kochi and 26 in Panjim. 101 transects in Kochi and 20 in Panjim had no trees. 84 trees in Kochi (36.05%) and 329 trees in Panjim (48.53%) out of the total observed, had cement, concrete or some other built structure around its base. Trees were identified up to the species level whenever possible and a total of 87 species were identified in Panjim (total city Simpson index =0.05; Shannon index=3.67) and 46 in Kochi (total Simpson index=0.05; Shannon index=3.37) across 31 and 20 families respectively. In both cities, the number of trees, number of species and the basal area of trees in a transect increased with road width (Fig 2).

**Table 1:**
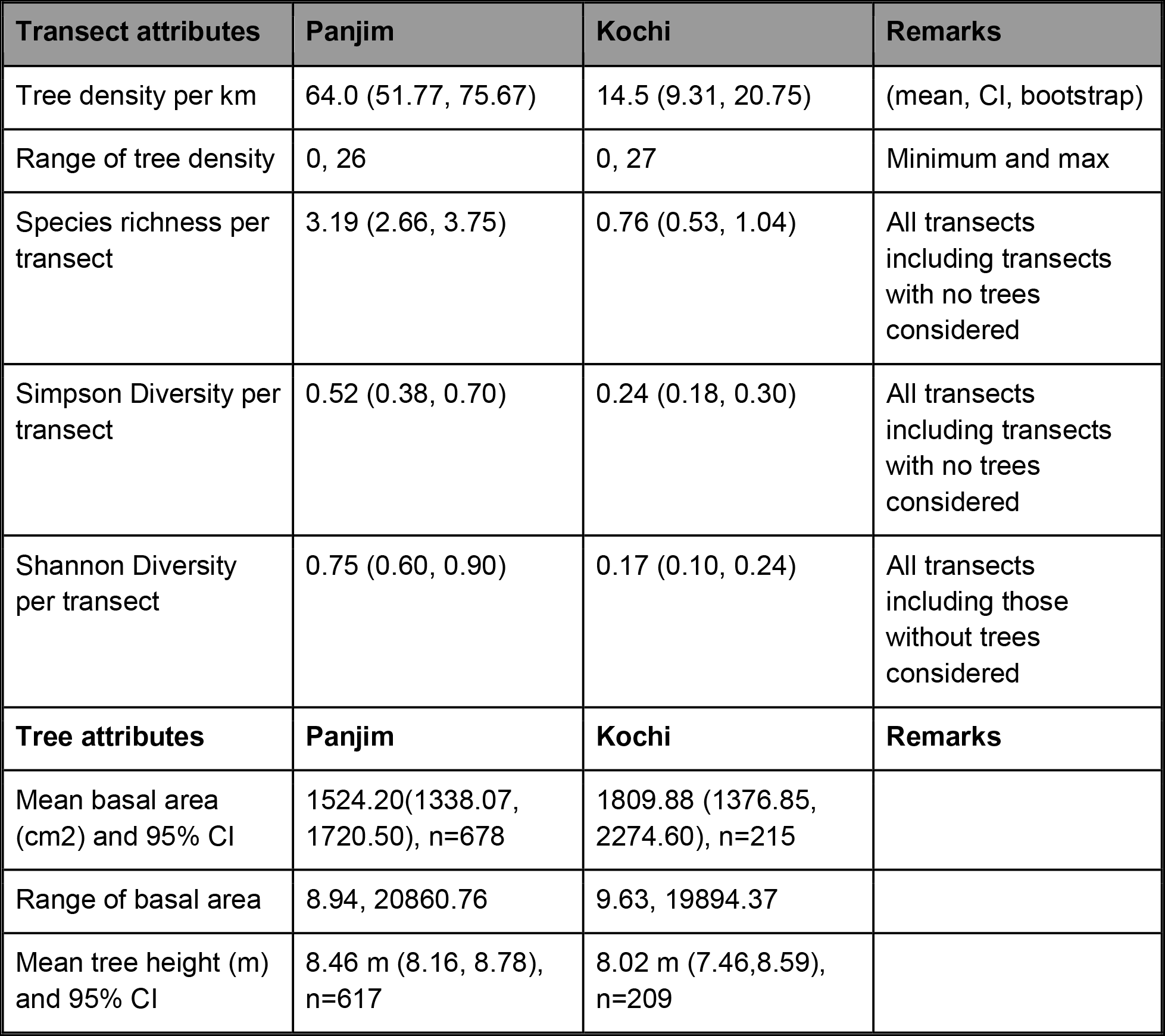
Summaries of transect and tree attributes observed across the cities. Values represent bootstrapped (n=9999) mean and 95% confidence intervals. Transect attributes reported for all transects in each city, except for Shannon diversity, reported only for transects with at least one tree. Tree attributes reported across all trees measured in each city, for which gbh and height values were reliably measured.

| **Transect attributes** | **Panjim** | **Kochi** | **Remarks** |
| --- | --- | --- | --- |
| Tree density per km | 64.0 (51.77, 75.67) | 14.5 (9.31, 20.75) | (mean, CI, bootstrap) |
| Range of tree density | 0, 26 | 0, 27 | Minimum and max |
| Species richness per transect | 3.19 (2.66, 3.75) | 0.76 (0.53, 1.04) | All transects including transects with no trees considered |
| Simpson Diversity per transect | 0.52 (0.38, 0.70) | 0.24 (0.18, 0.30) | All transects including transects with no trees considered |
| Shannon Diversity per transect | 0.75 (0.60, 0.90) | 0.17 (0.10, 0.24) | All transects including those without trees considered |
| **Tree attributes** | **Panjim** | **Kochi** | **Remarks** |
| Mean basal area (cm<sup>2</sup>) and 95% CI | 1524.20(1338.07, 1720.50), n=678 | 1809.88 (1376.85, 2274.60), n=215 |  |
| Range of basal area | 8.94, 20860.76 | 9.63, 19894.37 |  |
| Mean tree height (m) and 95% CI | 8.46 m (8.16, 8.78), n=617 | 8.02 m (7.46,8.59), n=209 |  |

**Figure 1:**
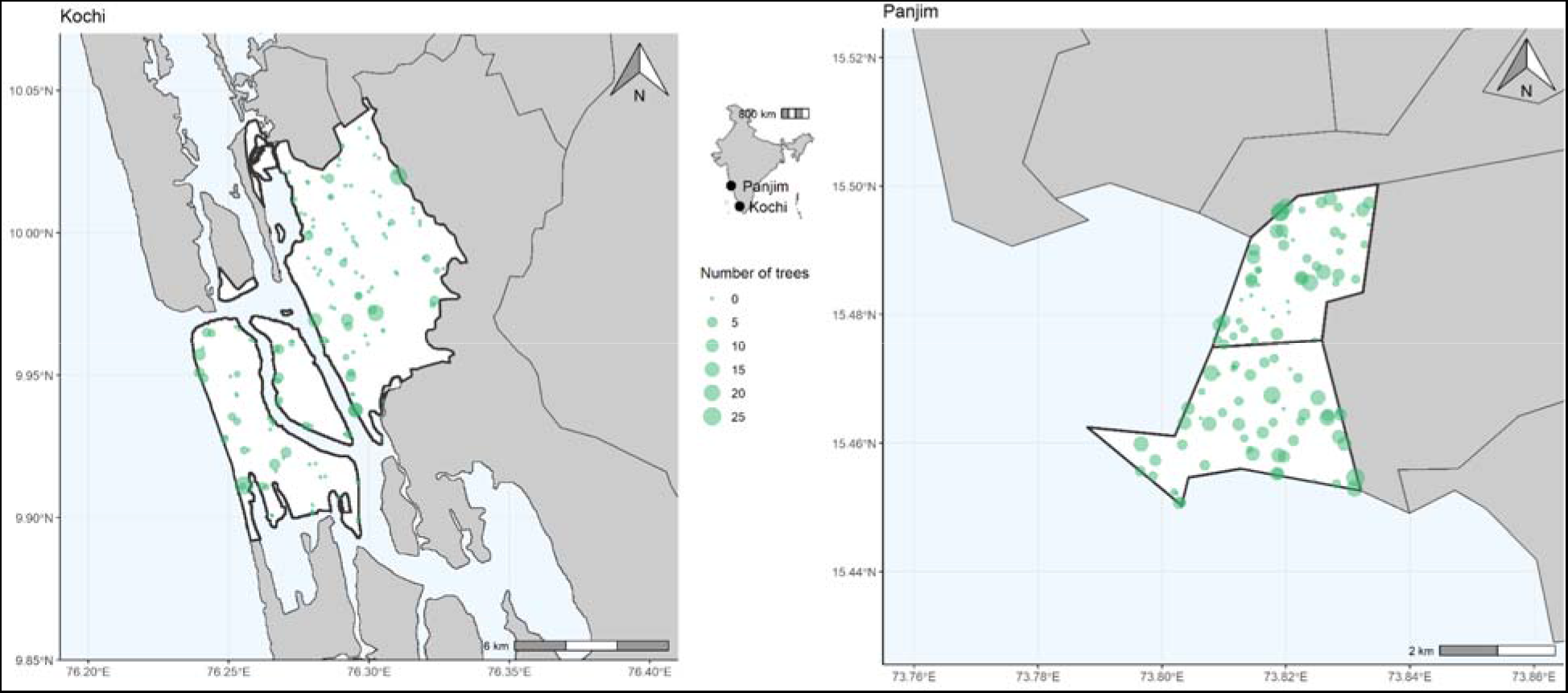
Study area map of a) Kochi and b) Panaji and Taleigao - together “Panjim”. White polygons represent administrative boundaries of the two cities considered and the green dots represent start points of road transects sampled during this study. The size of the green dots represents the abundance of trees in each transect. In the inset is a map of India with the locations of the two cities, Kochi and Panjim.

**Figure 2:**
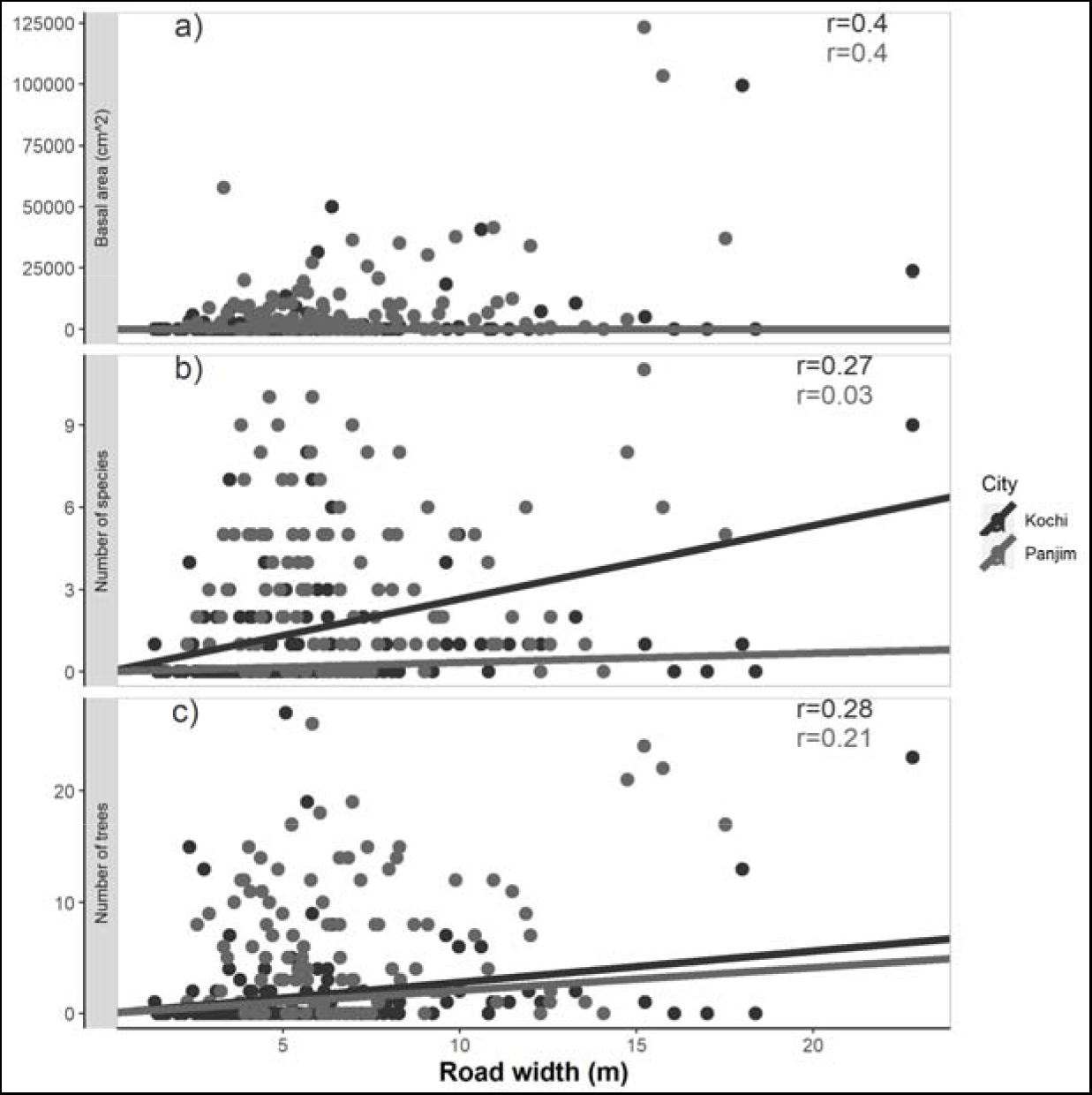
Tree community characteristics with road size in both Kochi and Panjim. a) Basal area of trees, b) number of species and c) number of trees as a function of road width. On the top right corner are Pearson correlation coefficients for each variable with road width in each city.

Observed trees in Kochi and Panjim were comparable in their basal area, height and dominant species identities. The density of tree basal area along roads was higher in Panjim (1.45×10^−3^ m^2^ per m^2^ of land) than Kochi (4.66×10^−4^ m^2^ per m^2^ of land). However, average basal area and height of observed trees in both cities were not significantly different, as seen by overlapping confidence intervals (Table 1).The maximum observed basal area and height were slightly larger in Panjim than in Kochi. Moreover, the three most commonly observed species in both cities are *Cocos nucifera*, *Albizia saman* and *Peltophorum pterocarpum*, ranked 1, 2 and 3 in Kochi and 2, 3, 1 in Panjim (Table 2)*. A. saman* (Rain tree) had the highest species importance value in both cities, followed closely by *P. pterocarpum* (Table 2). The ten most commonly observed species make up 86.7% and 76.2% of the total observed basal area in Kochi and Panjim respectively.

**Table 2:** Dominant species observed in Kochi and Panjim. Accepted name, common and local language name where available, number of trees, relative abundance, relative dominance, number of transects in which it was observed and species importance (relative abundance + relative dominance). Accepted name of each species is reported as per The Plant List, accessed using the R package *Taxonstand*. Tree species in each city are ordered in decreasing order of observed abundance and where abundance is equal, in decreasing order of importance.

| Accepted Species name | Common Name | Family | Total abundance | Relative abundance (%) | Total basal area (m<sup>2</sup>) | Relative dominance (%) | Frequency of occurrence | Relative frequency | Species importance |
| --- | --- | --- | --- | --- | --- | --- | --- | --- | --- |
| **Kochi** |  |  |  |  |  |  |  |  |  |
| *Cocos nucifera* | Coconut/then gu | Arecaceae | 29 | 12.39 % | 1.20 | 3.06% | 7 | 6.03% | 21.49 |
| *Albizia saman* | Rain tree | Fabaceae | 23 | 9.83% | 17.43 | 44.57 % | 12 | 10.34 % | 64.75 |
| *Peltophorum pterocarpum* | Copperpod tree | Fabaceae | 17 | 7.26% | 11.06 | 28.28 % | 3 | 2.59% | 38.13 |
| *Delonix regia* | Gulmohur/Va ka | Fabaceae | 14 | 5.98% | 1.53 | 3.92% | 2 | 1.72% | 11.62 |
| *Thespesia populnea* | Indian tulip/paruthy | Malvaceae | 13 | 5.56% | 0.51 | 1.29% | 1 | 0.86% | 7.71 |
| *Cassia fistula* | Golden showe tree/Kanikkonna | Fabaceae | 12 | 5.13% | 0.34 | 0.87% | 3 | 2.59% | 8.59 |
| *Wodyetia bifurcata* | Foxtail palm | Arecaceae | 12 | 5.13% | 0.28 | 0.72% | 1 | 0.86% | 6.71 |
| *Terminalia catappa* | Indian almond/Thallithenga | Combretaceae | 10 | 4.27% | 1.12 | 2.86% | 6 | 5.17% | 12.30 |
| *Mangifera* | Mango/Maav | Anacardiaceae | 7 | 2.99% | 0.53 | 1.34% | 6 | 5.17% | 9.51 |
| *indica* | u | eae |  |  |  |  |  |  |  |
| *Lagerstroemia speciosa* | Pride of India/Manimarruthu | Lythraceae | 7 | 2.99% | 0.34 | 0.88% | 4 | 0.34% | 7.32 |
| *Gliricidia sepium* | Sheemakkonna | Fabaceae | 7 | 2.99% | 0.11 | 0.27% | 4 | 0.11% | 6.72 |
| **Panjim** |  |  |  |  |  |  |  |  |  |
| *Peltopodium pterocarpum* | Copper pod tree | Fabaceae | 75 | 11.05% | 20.34 | 19.68% | 20 | 6.02% | 36.75 |
| *Cocos nucifera* | Coconut/Maad | Arecaceae | 62 | 9.13% | 3.76 | 3.64% | 21 | 6.32% | 19.09 |
| *Albizia saman* | Rain tree | Fabaceae | 59 | 8.70% | 30.63 | 29.64% | 24 | 7.23% | 45.56 |
| *Terminalia catappa* | Indian almond/Bodom | Combretaceae | 51 | 7.51% | 4.00 | 3.87% | 23 | 6.93% | 18.31 |
| *Mangifera indica* | Mango/Ambo | Anacardiaceae | 30 | 4.42% | 9.95 | 9.62% | 17 | 5.12% | 19.16 |
| *Delonix regia* | Gulmohar | Fabaceae | 25 | 3.68% | 2.48 | 2.40% | 8 | 2.41% | 8.49 |
| *Bombax ceiba* | Red silk-cotton/Sanvor | Malvaceae | 24 | 3.53% | 1.58 | 1.53% | 11 | 3.31% | 8.37 |
| *Polyalthia longifolia* | False Ashoka/Ashok | Annonaceae | 23 | 3.39% | 3.48 | 3.37% | 5 | 1.51% | 8.26 |
| *Lannea coromandelica* | Indian ash tree/Moi | Anacardiaceae | 20 | 2.94% | 1.65 | 1.60% | 10 | 3.01% | 7.56 |
| *Cassia fistula* | Indian laburnum/Amaltas | Fabaceae | 16 | 2.36% | 0.86 | 0.83% | 8 | 2.41% | 5.60 |

### Human-tree relationships from our interview of residents

Overall, respondents in both cities reported a net decline in tree cover in the recent past. Aggregating across responses from all residents who have lived in the city for 10 years or longer, respondents in Kochi on average reported a 28.35% decline in tree abundance in the recent past (95% CI: −30.49%, −26.06%) while respondents in Panjim reported a 11.65% decline (95% CI: −14.23%, −9.10%) (Fig 3a). Residents in Kochi perceived a larger decline in tree abundance during their stay in the city than residents of Panjim. Respondents in these cities reported associations with green spaces and visited parks. 72 respondents in Panjim (39.34%) and 105 in Kochi (33.43%) reported having visited a park or a green space within a year or less. Out of this, 24.59% in Panjim and 9.23% in Kochi visited within the month or less and 13.66% and 2.86% respectively visited within the week. Respondents in Kochi were most likely to report a 25 to 50% decline (68.33% of 259 respondents), while in Panjim a 5 to 25% decline in tree cover was the most likely response (38.13% of 160 respondents). 67 respondents in Kochi (21.33%) and 22 respondents in Panjim (12.02%) reported that trees were frequently cut down in their areas for various purposes while 154 Kochi respondents (40.04%) and 114 Panjim respondents (62.3%) reported that trees were rarely or never cut down in their area. Land clearance for buildings was the most commonly quoted reason for cutting down trees frequently in both cities, much larger than land clearance for roads, footpaths, amenities or to mitigate risk to life and property (Fig 3b). On the other hand, 5.10% and 37.70% of respondents in Kochi and Panjim respectively reported that trees are frequently planted in their area, while 59.24% and 37.70% said that trees were rarely or never planted in their area.

**Figure 3:**
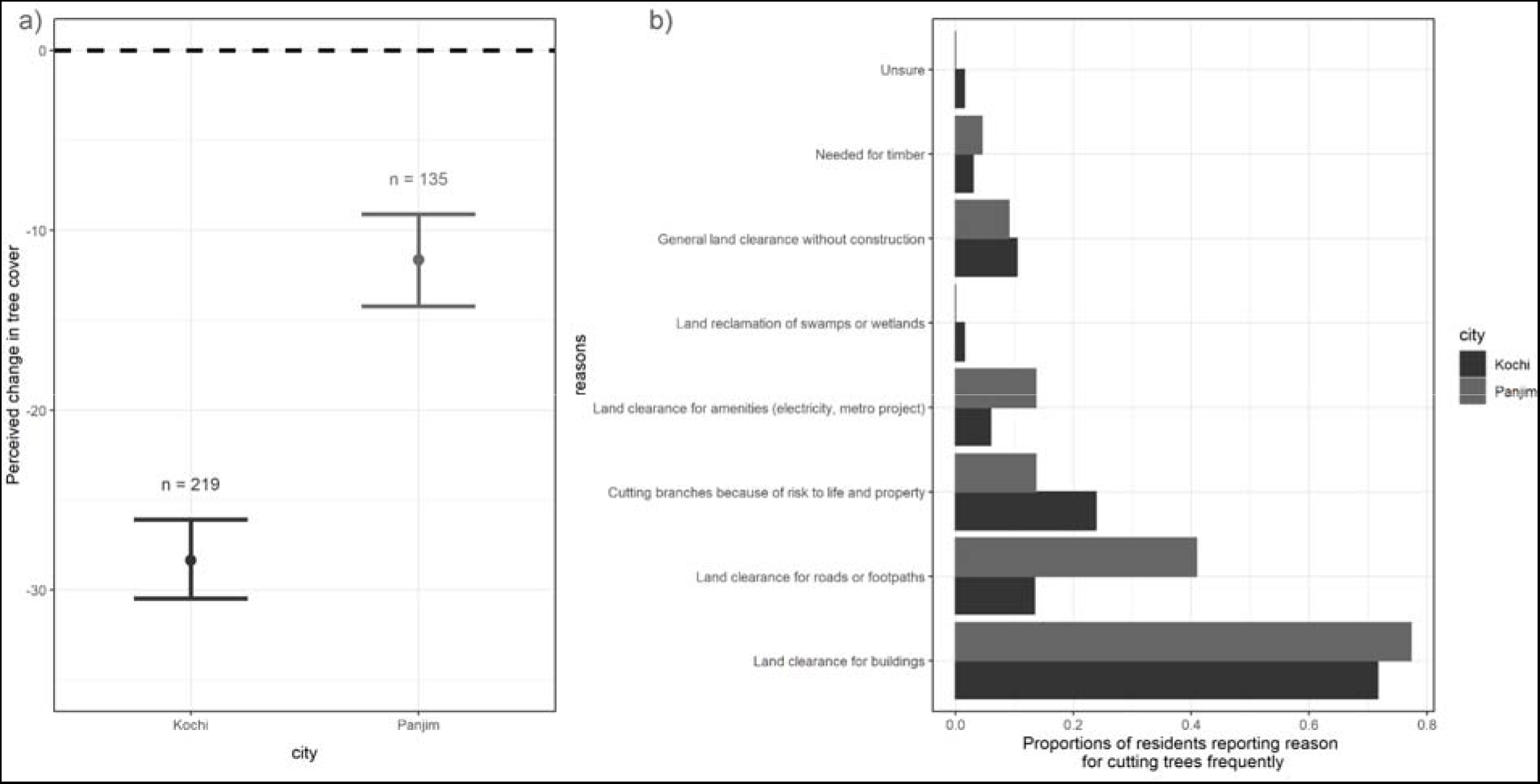
Perceptions of change in Kochi and Panjim among long-term residents (10 years or longer). a) Perceived change in tree cover in the two cities; the dot represents the bootstrapped mean change and error bars represent 95% CI (bootstrap n=9999). Sample sizes reported above each city. Dotted line is the zero line. b) Reported reasons for frequently cutting down trees and the proportion of residents reporting reasons in each city.

Trees in both these cities were reported to provide various provisioning, regulating and cultural services (Table 3). Correspondingly, high values of regulating services were most often reported in both cities, while direct provisioning services were least reported. For both provisioning and cultural services, respondents in Panjim were significantly more likely to report high value (Fig 4). Respondents in both Kochi and Panjim reported the high value of trees for shade and water supply. Almost all respondents found trees beautiful. Respondents also noted the religious values of trees as well as their presence next to prominent buildings, but these values were larger in Panjim than in Kochi. In Panjim, trees were also reported to be more used and valued for consumption and medicine, while trees in Kochi were almost never used for medicine, and a substantial proportion of respondents reported no consumptive value (Table 3). Significantly higher proportion of respondents reported no value for specific services in Kochi for food, medicine, boundary marking and as associated with cultural and important buildings (Table 3). High reported value on one service was often positively correlated with value of other services, more so for Panjim than for Kochi (Fig 4 b, c). Moreover, these correlations were not limited to uses within ecosystem service categories; the number of functions reported as high value under each category correlated with value in other categories (Fig 4). The correlation between provisioning and cultural service value in Panjim was the highest across all pairwise correlations.

**Table 3:**
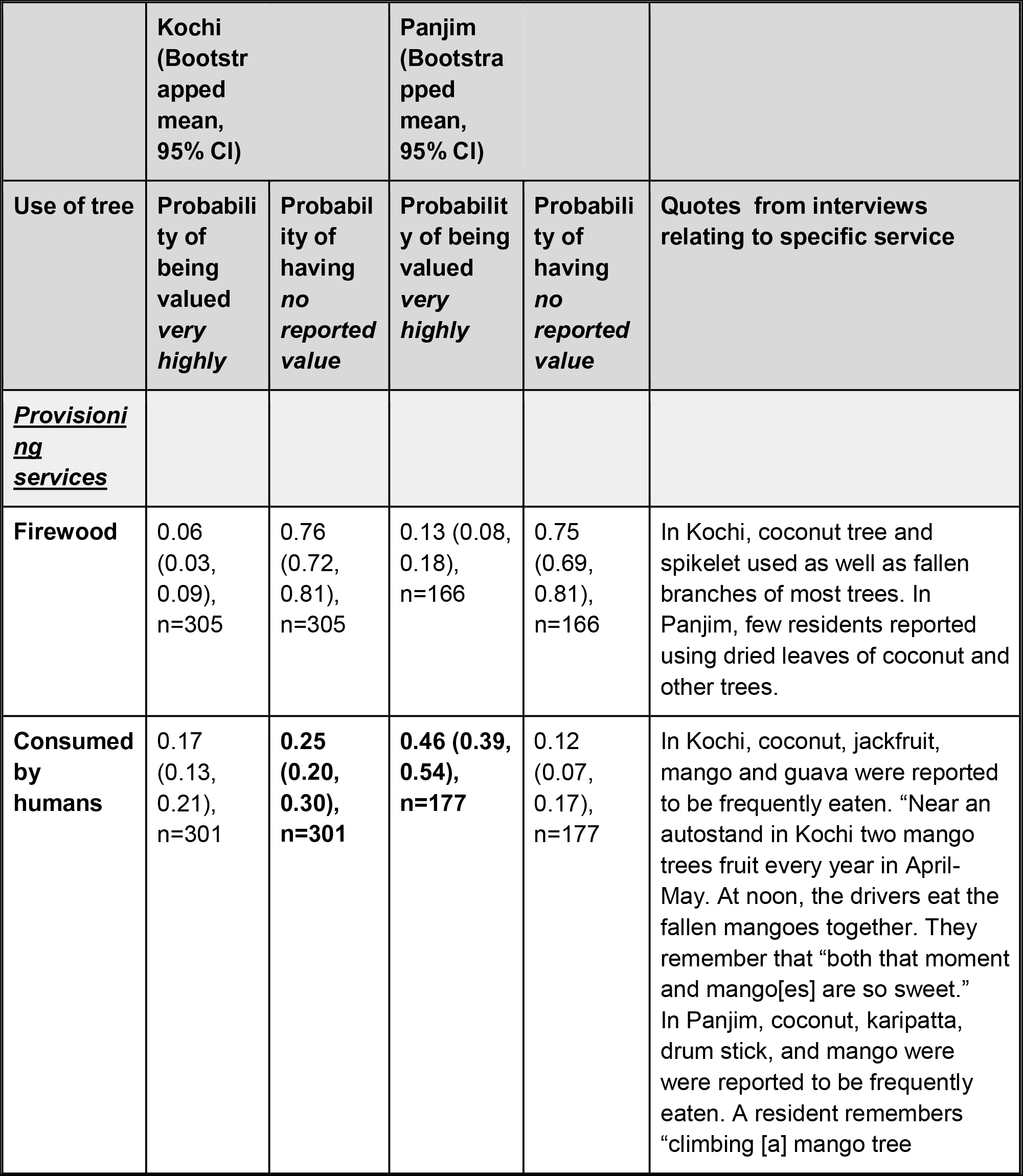
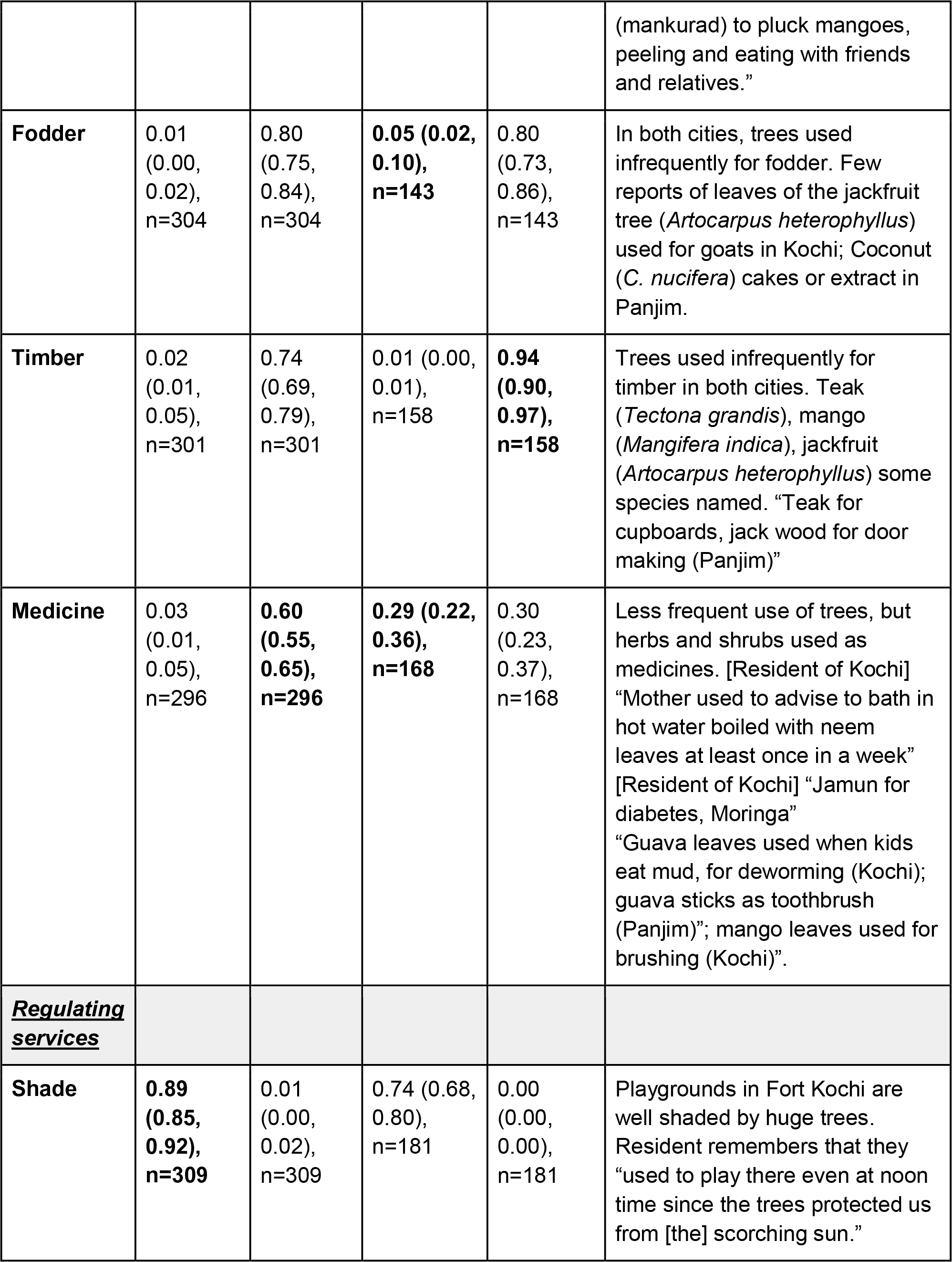
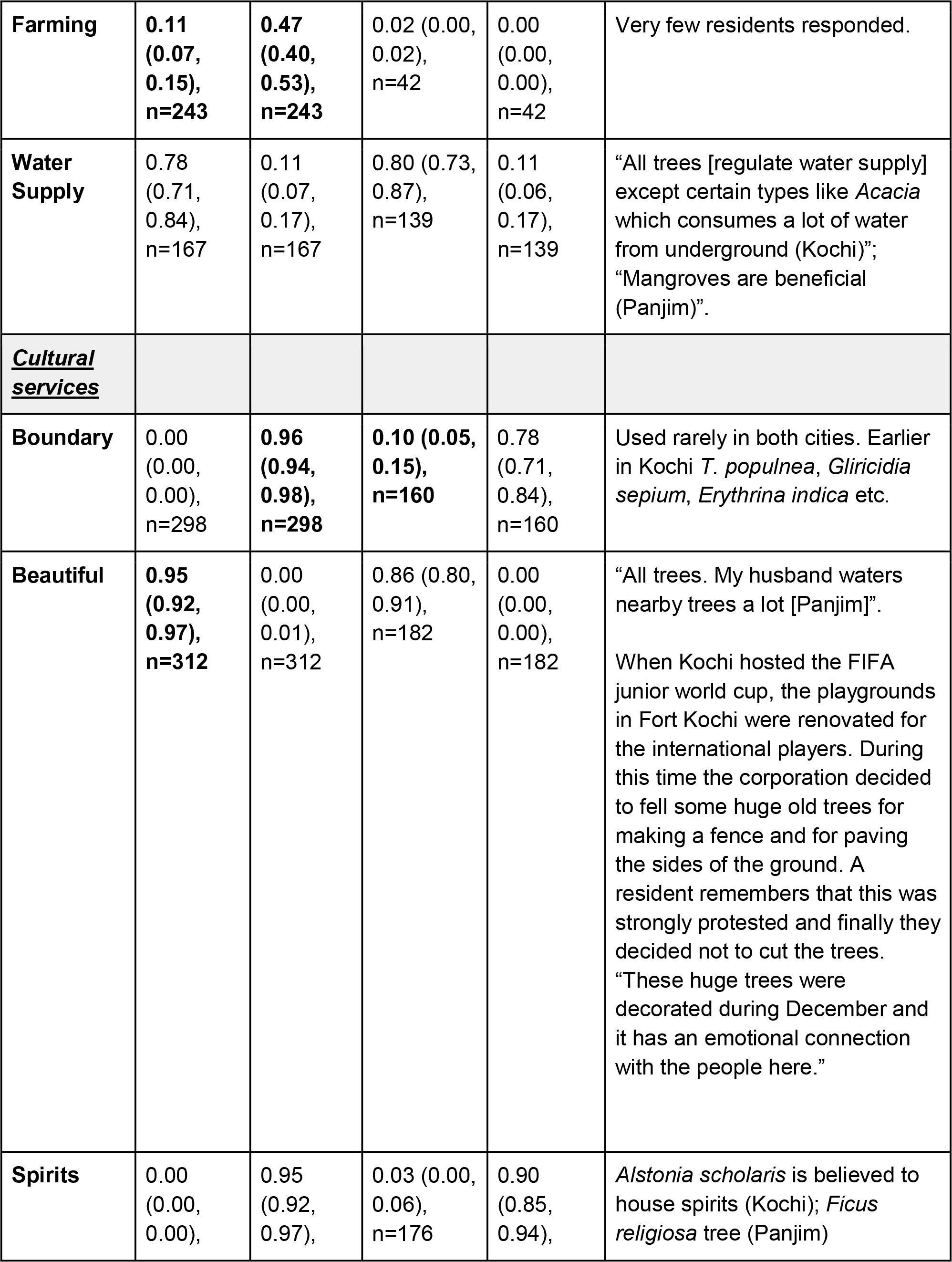
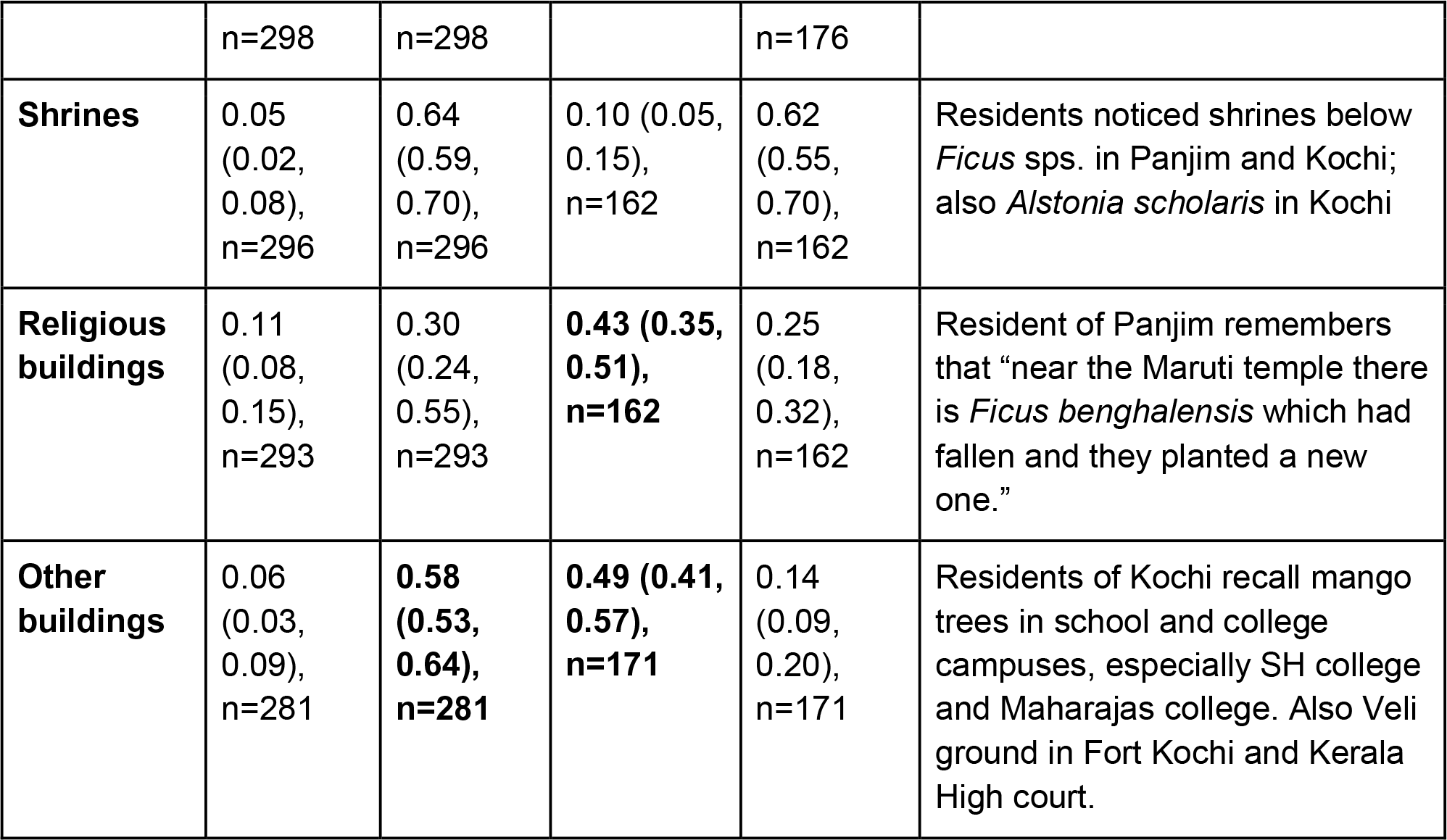
Perceived ecosystem services from street trees in Kochi and Panjim. For each service in “provisioning”, “regulating”, or “cultural” ecosystem service category, we report bootstrapped mean and confidence intervals of the proportion of respondents who reported *very high value* or *no value* in their area. Where uses are significantly different in the two cities, with no overlaps in confidence intervals, the higher value is highlighted in bold.

**Figure 4:**
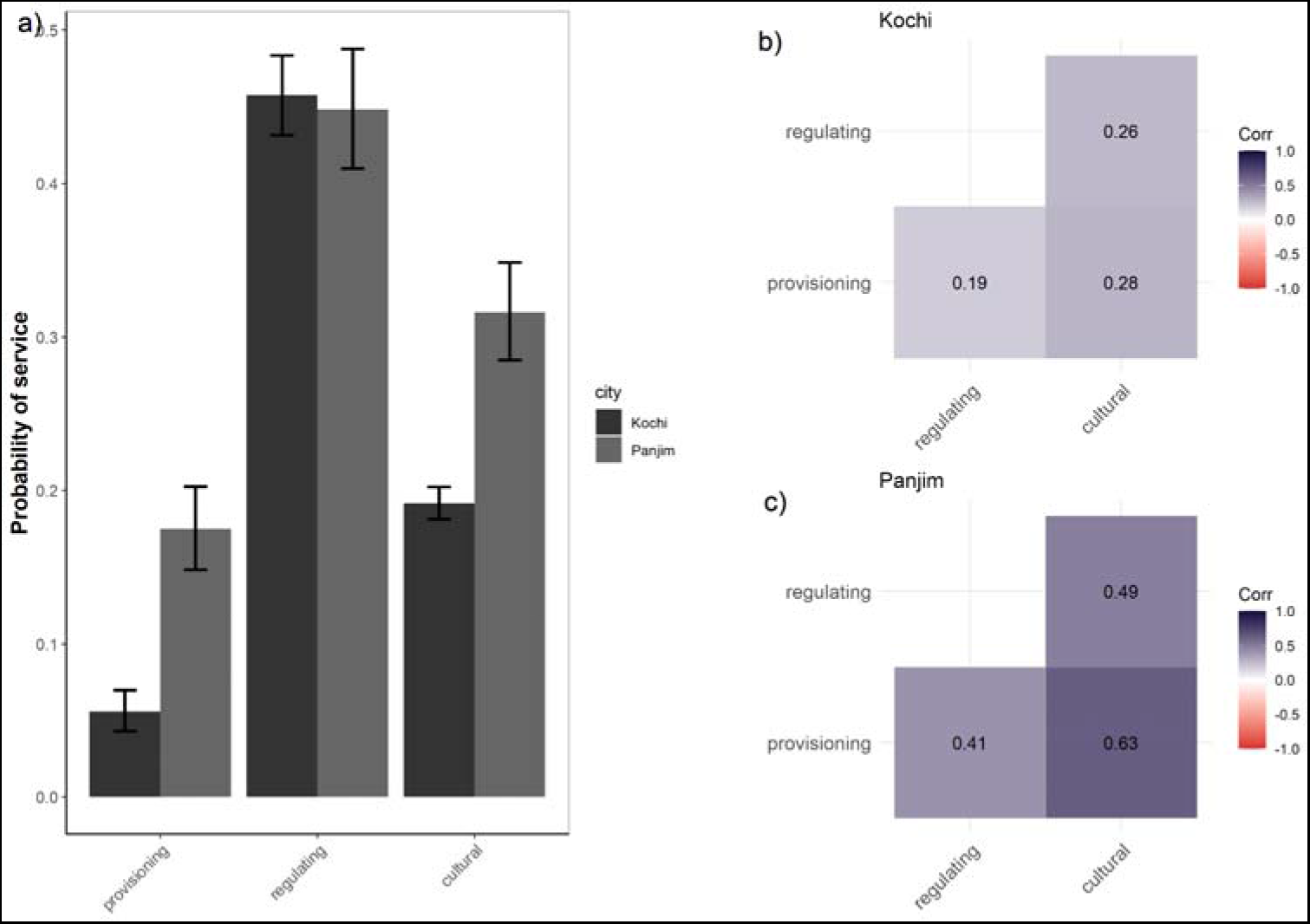
Ecosystem services provided by trees in each city. a) The proportion of the three types of service provided by trees in each city. The height of the bar represents the mean proportion and error bars represent 95% CI (bootstrap n=9999). Correlations between the number of regulating, provisioning and cultural services reported having “very high value” in the area in b) Kochi and c) Panjim.

## Discussion

The global focus of urban planning for sustainability has been on metropolises. In India, one of the fastest growing economies in the world, intermediate cities and census towns have a growing influence, but their opportunities and challenges and their potential contribution to understanding global urban sustainability remain unexplored (Tripathi & Mitra, 2022). In this study, we conducted the first systematic documentation of the socio-ecological systems of two growing coastal cities in India, Kochi and Panjim, known for their natural beauty and built heritage. We documented low density of street trees and a reported decline in tree cover in the recent past. Despite urbanization being fairly recent in these cities, residents infrequently value trees for provisioning services like food, medicine and fodder. However, we documented high cultural value for trees and strong, place-based narratives of the importance of trees. While the rapid scale and change of planetary urbanization has gained much attention, combining this top-down approach with understanding of socio-spatial contexts of places is imperative and requires further attention (Kumar & Shaw, 2020).

In Kochi and Panjim, two small urban centers with a growing trend in urbanization, we found evidence of net decline in tree cover in the recent past and corresponding low street tree density. Presently, mean tree cover along roadsides is lower than many other Asian countries and values in Kochi are lower than Bengaluru as reported in Nagendra & Gopal, 2010 (mean across all roads for Panjim = 64 per km of road and Kochi = 14 compared to Bengaluru mean for medium roads = 73.0, narrow roads = 52.1). Some of these differences could be due to the difference in size and scale of infrastructure in these cities (Kuriakose & Philip, 2021). The core of Panjim city has a planning history of at least 175 years with a grid-wise plan laid out, whereas beyond a few streets, Kochi’s growth has mainly involved unplanned urban sprawl (Abhimanyu et al., 2020). According to news reports, over 1 lakh trees in public spaces in Kochi have been cut during the last 9 years for several expansion projects and replanting efforts have not kept up with this loss (Deccan Chronicle, 2019). Our interviews reflect this change; residents in Kochi reported 28% decline in tree cover on average, while in Panjim, reported decline was smaller in scale, at 11% (Fig 3a). Residents in both cities quoted land clearance for buildings as the most frequent reason for cutting trees in their area (Fig 3b), in alignment with urbanization trends in metropolises as well. Many smaller roads in Kochi did not have pavements, with walls of private property directly marking the road edge, leaving no possibility for planting trees, as can be noticed by the number of zero-tree transects in Fig 2.

The tree community in these cities represented a mix of exotic avenue species and species native to these coasts. *A. saman* and *P. pterocarpum* were the two most important species recorded among the street trees of these cities, and the family Fabaceae, of which these species are members, made up more than 50% of the basal area in both cities, and more than 75% in Kochi. These two species commonly observed in our study are also common avenue species across cities in India; they make up the most abundant species along larger roads in Bengaluru (Nagendra & Gopal, 2010). However, being coastal, coconut, *C. nucifera* was also an abundant tree species in both cities (Table 2). Moreover, deviating further from inland cities previously studied, we found that a few coastal littoral species that are part of the native vegetation also occurred commonly - *Thespesia populnea* in Kochi and *Terminalia catappa* in Panjim made up important parts of the sampled tree community (Table 2). The persistence of these native species in the local community could potentially influence which ecosystem services residents value.

In these small urban centers, residents were more likely to value regulating services of trees, compared to direct benefits like food and medicine, but strong cultural values were also notable. Trees were used for a variety of purposes in these cities, but residents of Panjim had higher frequencies of use and trees were seen as more multifunctional (Fig 4). Substantial proportion of respondents within both cities visited parks within a year or less, and residents of both cities concur on the value of trees for providing shade, shelter and for maintaining the water table. Notably, in both cities, there was agreement on the aesthetic value of trees (Table 3). Qualitative information from interviews also revealed strong human-tree relationships of many residents, through cultural and personal memories of trees (Table 3). Many native species were recounted in these accounts; for example, residents reported the use of T. populnea as live boundaries in Kochi, a practice that was documented in the tree surveys as well. These intangible benefits of trees ranked high, suggesting high inherent cultural and social value for trees and human connection for nature in these cities (Gopal et al., 2018; Graça et al., 2018; Jaganmohan et al., 2018; Jennings et al., 2016). The indirect benefits of trees in these cities could be linked to the tree community assessed; ornamental and avenue trees make up a large proportion of the street-side and publicly accessible trees (Kishen, 2005). Fruit trees and medicinal trees make up a much smaller composition of the tree community, unlike in Bengaluru where species like neem (*Azadirachta indica*), valued for medicinal use, and jamun (*Syzygium cumini*), a fruit tree, are common roadside trees, providing direct benefits to residents (Mundoli et al., 2017; Nagendra & Gopal, 2010). Cultural values of trees among residents are also evident in civic engagement in nature-based citizen science as well as activism in these cities. Platforms such as eBird and SeasonWatch have the highest records from Kerala; a citizen-led project, the Kerala Bird Atlas, recently published Asia’s most extensive bird atlas (Praveen, 2022). In Goa, environmental citizenship related to coal and Goa’s largest protected areas (Jamal, 2020; Lobo, 2020) also suggests the potential to harness this stewardship towards large-scale biophilic planning.

Our study assessed only street trees, but preserving other green spaces in cities like heritage trees, urban forests and monitoring their health will be imperative in the coming decades as these cities continue to grow. Large-sized trees, and species that attain large sizes, like *Ficus* sp. and *A. saman* are keystone species for urban fauna as well as contribute disproportionately to ameliorating urban heat island effects (Sinha et al., 2021; Vailshery et al., 2013). Expansion and tree cutting in Panjim have focussed on large sized species (P. Fernandes, n.d.), potentially leading to the loss of multiple services simultaneously. Moreover, street trees represent a small proportion of the green spaces in these cities; private gardens and private fruiting trees in these cities are anecdotally more utilized, potentially influencing human-nature relationships. However, as urbanization increases in pace, home gardens and backyards within the city are likely to decrease, as observed in many metropolises especially in the developing world, necessitating investment into biophilic planning (Dobbs et al., 2018; du Toit et al., 2018; Imam & Banerjee, 2016). Systematic, context-based ecological information on these other dimensions of urban greenery is necessary to add to our understanding of urban sustainability in the global South.

Further understanding of human-nature relationships in these growing cities could help fill gaps in our knowledge of urban sustainability in a rapidly developing global South context. India currently has 8 metropolises, but over 200 smaller urban centers, where we know little about growth and urban sustainability. Out of these, 100 cities are funded under the National Smart Cities Mission aimed at driving economic growth and quality of life. Incorporating diverse citizen goals, some cities in developed countries have developed and implemented robust biophilic management plans, highlighting the potential to pioneer such efforts in urbanizing cities in India as well (Kloster et al., 2021). Panjim and Kochi are notable in their high participation towards citizen science and environmental stewardship which could aid biophilic planning. However, the scope of biophilic research and planning in these cities needs to broaden to account for coastal protection, ecotourism and mental health. Expanding the scope of existing initiatives (whether the Serve to Preserve tree mapping initiative in Kochi or the Living Heritage tree health mapping mobile application in Goa) as well as expanding research across rapidly urbanizing cities in India is necessary to set baselines for understanding trajectories of change and to inspire resilient urban planning.

## Acknowledgements

We are grateful to the residents of both Kochi and Panjim. Authors KA and NV grew up in these cities respectively and continue to spend time each year. We are grateful to Kesia Ann, Anjal Saji, Krishnaja, Olivia Chandy and other research assistants who collected and entered the data in Kochi. We also thank Fr. Jose John and the administration at Sacred Hearts College, Thevara for logistical support. We are thankful to Siddhi Gawas, Shubha Kauthankar, Nikita Gawas, Monaliza Dias Sapeco, Charles Po, Ankita Gawas, Asmita Simepuruskar, Nita Kinlekar, Shruti Kauthankar for their data collection efforts and research assistance in Panjim. We thank Seema Mundoli for initial discussions and conceptualizing the study. We are also grateful to Azim Premji University for financial support.

