## Supplemental Table 1 for "Beyond the metropolis: Street tree densities and resident perceptions on ecosystem services in small urban centers in India"

**Supplementary Information**

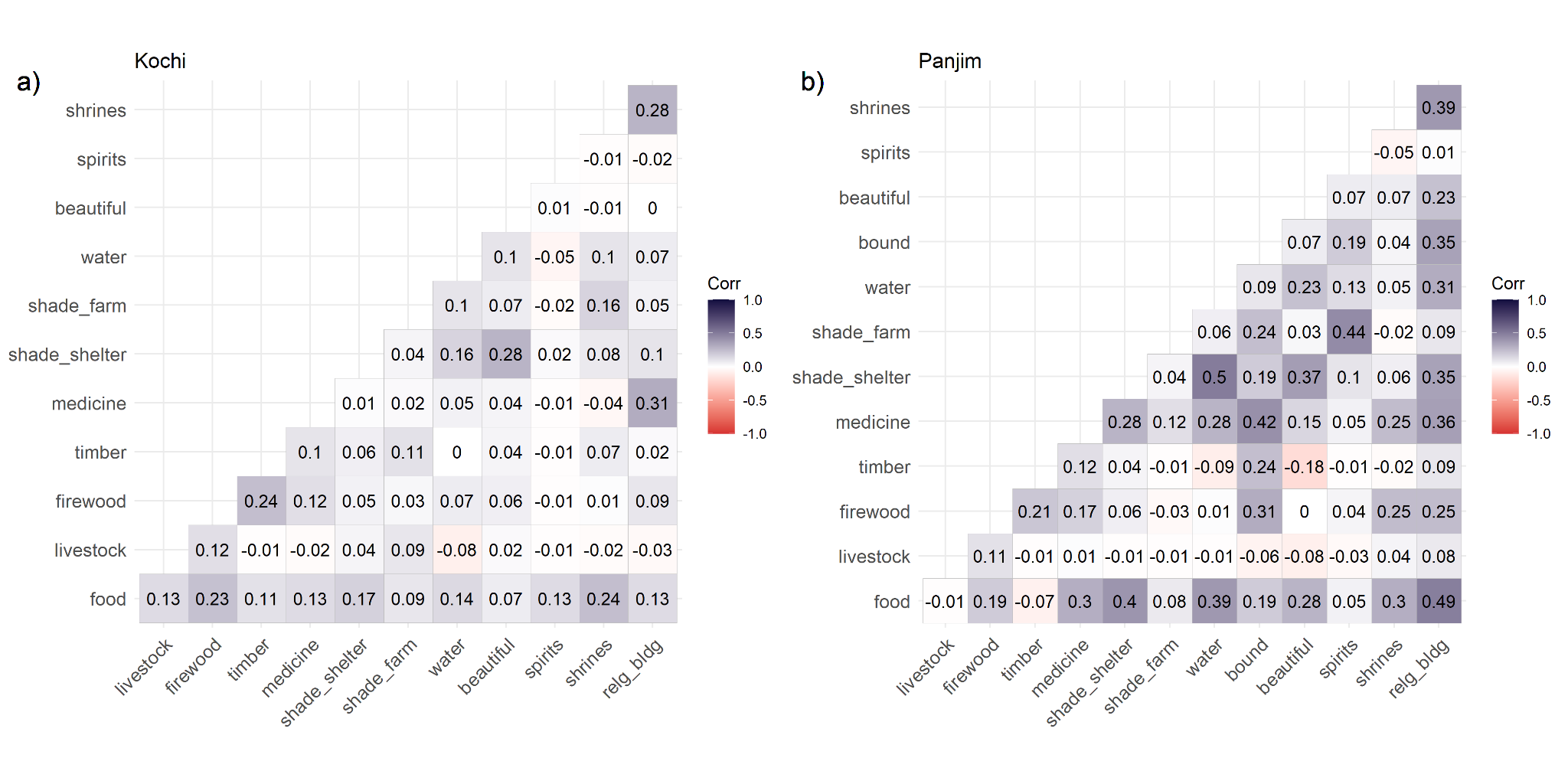

Figure S1: Ecosystem services provided by trees in each city. Correlation between reported “very high value” for each service from interviews of residents in a) Kochi and b) Panjim.
